# Contextual effects on binocular matching are evident in primary visual cortex

**DOI:** 10.1101/541466

**Authors:** Reuben Rideaux, Andrew E Welchman

## Abstract

Global context can dramatically influence local visual perception. This phenomenon is well-documented for monocular features, e.g., the Kanizsa triangle. It has been demonstrated for binocular matching: the disambiguation of the Wallpaper Illusion (Brewster, 1844) via the luminance of the background (Anderson & Nakayama, 1994). For monocular features, there is evidence that global context can influence neuronal responses as early as V1 (Muckli et al., 2015). However, for binocular matching, the activity in this area of the visual cortex is thought to represent local processing, suggesting that the influence of global context may occur at later stages of cortical processing. Here we sought to test if binocular matching is influenced by contextual effects in V1, using fMRI to measure brain activity while participants viewed perceptually ambiguous “wallpaper” stereograms whose depth was disambiguated by the luminance of the surrounding region. We localized voxels in V1 corresponding to the ambiguous region of the pattern, i.e., where the signal received from the eyes was not predictive of depth, and despite the ambiguity of the input signal, using multi-voxel pattern analysis we were able to reliably decode perceived (near/far) depth from the activity of these voxels. These findings indicate that stereoscopic related neural activity is influenced by global context as early as V1.

## INTRODUCTION

The horizontal offset between the two eyes provides the basis for binocular disparity signals, which are a powerful cue for inferring the three-dimensional structure of the surrounding environment. Binocular neurons in V1 receive feedforward input from monocular neurons in the lateral geniculate nucleus (Barlow, Blakemore, & Pettigrew, 1967; Hubel & Wiesel, 1970). By computing the retinal disparity between corresponding elements in a scene, the brain estimates the sign and magnitude of their depth relative to fixation. However, while local disparity signals may provide an initial estimate of depth, we know that perception is also influenced by global contextual signals. For instance, two objects of equal luminance can appear to have different levels of brightness if they are observed in different contexts (**Figure 1a**). Similarly, the perception of luminance defined contours can be induced by the presence of spatially distinct elements within a visual scene if they are arranged such that the visual system extrapolates the presence of an occluding surface, as in the Kanizsa triangle illusion (Kanizsa, 1979; **Figure 1b**).

**Figure 1.**
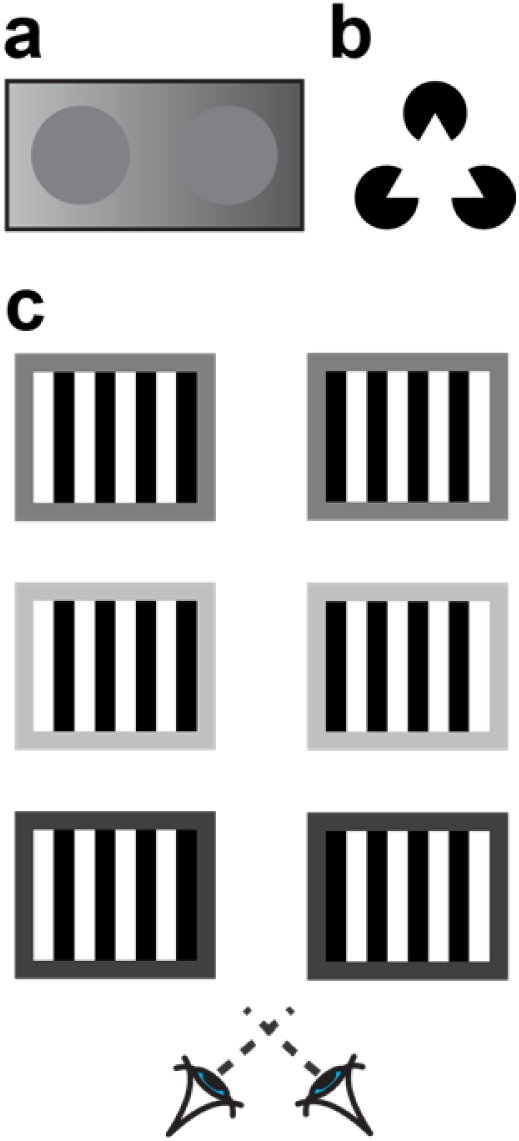
Demonstrations of spatial contextual effects. **a)** Simultaneous brightness contrast, the perceived luminance of the disks is determined by the luminance of the background. **b)** A Kanisza triangle, the ‘pacman’ disks induce illusory luminance contours to complete the perceptually inferred triangle. **c) Stereograms of the Wallpaper stimulus designed for cross-eyed fusion.** When fusing the left and right eye images in the top pair, the reader should experience and ambiguous perceptual interpretation that alternates between the central region appearing ‘near’ or ‘far’ relative to the surround. Changing the luminance of the background from mid-grey to light grey (middle) or dark grey (bottom) changes the perceptual interpretation: the middle stimulus should look ‘far’ while the bottom stimulus should appear ‘near’.

Spatial context can influence the perception of depth. For instance, a repetitive pattern presented to the two eyes can pose a challenge to the visual system in that a range of different interpretations of the matches between the left and right eyes’ views are possible – the Wallpaper Illusion (Brewster, 1844). Spatial context can influence which interpretation is selected by determining which of the similar elements are matched between the two eyes. In particular, when the left and right eyes are presented with square-wave gratings with a 180° phase offset (such that a light bar in the left eye corresponds to a dark bar in the right eye and vice versa) there are two ways that the images can be stereoscopically matched. The images can be “shifted” nasally or temporally to align the stripes of the in the centre of the gratings while producing half-occlusions at the edges. When the background luminance of the grating is mid-way between the light and dark bars, the nasal/temporal stereoscopic matching solutions are equally probable, producing a bistable situation in which the perceived depth of the grating alternates between near vs. far interpretations. By manipulating the background luminance of the grating, it is possible to constrain the perceptual interpretation of the stimulus from its ambiguous state so that one perceptual interpretation is chosen (Anderson & Nakayama, 1994; **Figure 1c**). This is thought to occur as a result of the visual system seeking to preserve contrast polarity during the matching of monocular inputs. Critically, while this matching constraint is only present at the left and right edges of the stereogram, the entire pattern is perceived as consistently near or far. This suggests a neural mechanism sensitive to extensive portions of the viewed scene.

It is interesting to consider how such displays might influence the responses of binocular neurons in the visual cortex. Within primary visual cortex, neurons have small receptive fields, meaning that they could only register the disparity of a small portion of the display. While neurons sampling the edges of the display may have their responses disambiguated by the contrast of the background, we would expect those sampling from the centre of the wallpaper pattern to provide ambiguous information as to whether the image features projected onto one retina should be matched nasally or temporally with respect to the image on the other retina (**Figure 2**). Disambiguation of the signal could occur as early as V1: lateral or feedback connections from neurons receiving input from the edges of the pattern could influence the local (ambiguous) activity of those receiving input from the centre. However, electrophysiological recordings in macaques suggest that the activity of V1 neurons may not be representative of perceived depth. Cumming and Parker (2000) manipulated the perceived depth of a stimulus whilst maintaining the same retinal input within the receptive field of binocular neurons in V1 and showed the activity of these neurons was largely unaffected by the manipulation. Further, Cumming and Parker (1997) showed that binocular neurons in V1 respond to anti-correlated random-dot stereograms, i.e., in which the polarity of dots is reversed between the left and right eyes, which do not produce a percept of depth. These results suggest that V1 activity does not always accord with the stereo depth percept of the observer, and thus that the influence of spatial context may occur at a later stage of processing.

**Figure 2.**
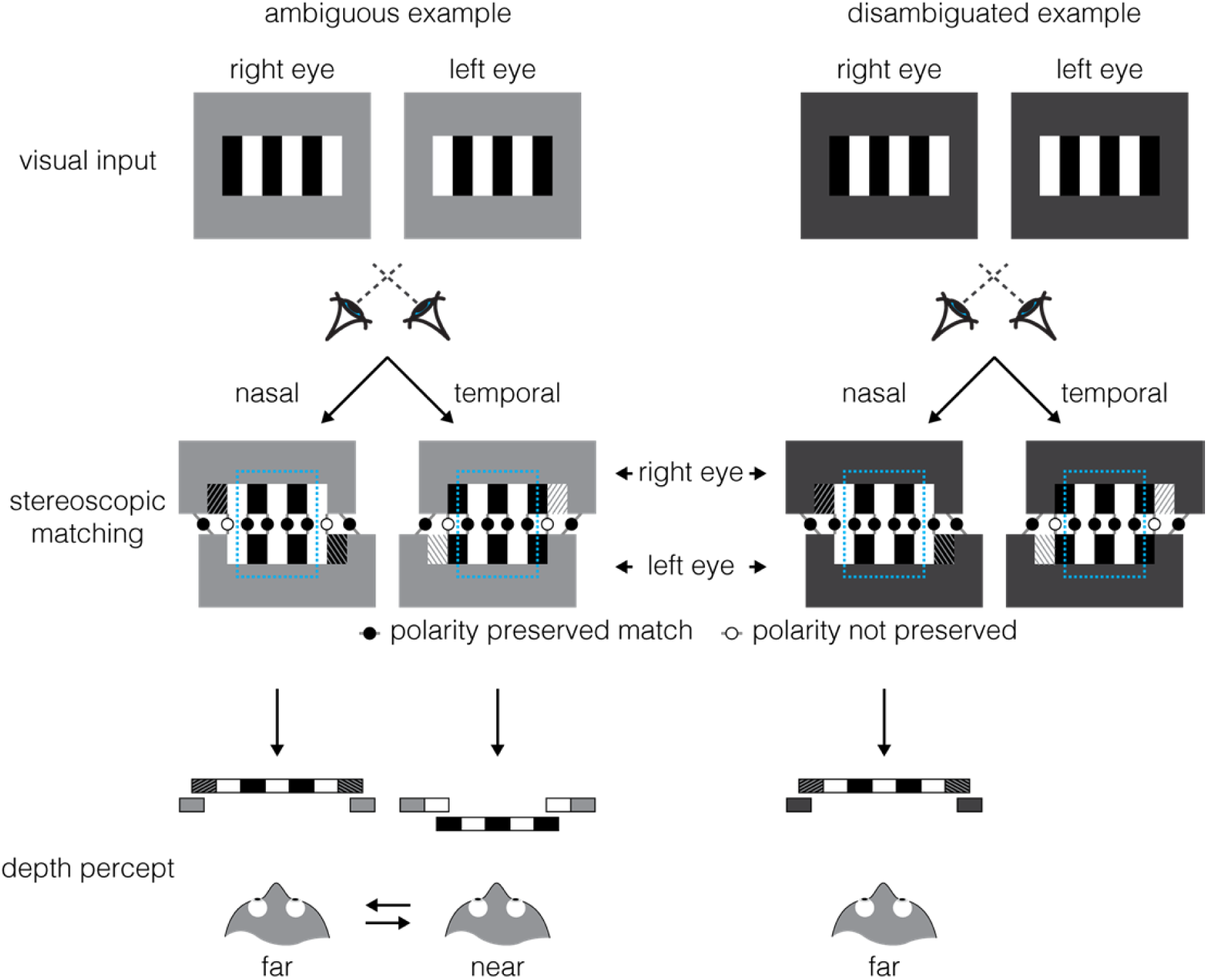
Illustration of contextual influence on binocular matching of an ambiguous ‘wallpaper’ image. The left and right eyes are presented with square-wave gratings, with a 180° phase offset. The images can be stereoscopically matched in two ways (i.e., nasal/temporal) with half-occlusions at the edges (oblique patterns: leftward tilt, viewed by left eye; rightward tilt, viewed by right eye). At the centre of the pattern (cyan rectangle) near vs. far matching solutions are equally likely, and thus ambiguous, as both preserve polarity contrast (solid circles). As shown in the (left-side) ambiguous example, if the luminance of the background is at the midpoint between the light and dark stripes of the pattern, this is also true for the edges, resulting in an ambiguous bistable (near/far) percept. However, as shown in the (right-side) disambiguated example, if the background is darker (or lighter) than the midpoint, matching solutions produce outcomes of different perceptual likelihood at the left and right edges of the pattern – while polarity contrast is preserved for the far solution, i.e., dark-light is matched with dark-light (solid circles), the near solution violates this rule, i.e., dark-dark is matched with dark-light (empty circles). In V1, receptive fields are relatively small and only sample from local regions of the pattern. While neurons with receptive fields positioned at the edges of the pattern receive a disambiguating signal, the majority of neurons (sampling from region within the cyan rectangle) receive ambiguous signal. Despite the disambiguating signals only featuring at the edges of the pattern, they are sufficient to produce a reliable depth percept over its entire surface.

Here we rely on the spatial resolution of fMRI to examine information about the responses to depth stimuli within the visual cortex. Specifically, in V1, where receptive fields are relatively small, neurons sampling information from the edges encode depth signals that are constrained by the local contrast signals. By contrast, neurons sampling from the centre of the pattern receive ambiguous input that can be matched in a temporal or nasal direction. If local activity in V1 is unaffected by the global context of the edges, the pattern of fMRI activity in a subregion of V1 responsive to the centre of the pattern should not be predictive of apparent depth. However, if perceived depth can be reliably predicted from activity in this region, this shows that contextual influence occurs as early as V1.

Thus, to examine whether global context influences fMRI activity in V1 during binocular matching, here we use multi-voxel pattern analysis (MVPA) of BOLD activity to decode depth from regions of V1 receiving ambiguous binocular input whilst perception is influenced by spatial context. We find some evidence that perceived depth can be decoded from regions of V1 that receive ambiguous depth information. This suggests a modulation of the local BOLD signal in V1 that is influenced by the global process of establishing binocular correspondence.

## MATERIALS AND METHODS

### Observers

Fifteen healthy observers from the University of Cambridge with normal or corrected-to-normal vision participated in the fMRI experiment. The mean age was 26 yr (range = 21–32 yr; 9 women, 5 men). Two participants’ data were excluded from the analysis because of excessive head movements during scanning, which prevented voxel correspondence necessary for multi-voxel pattern analysis (MVPA). Excessive head movement was defined as ≥1 mm displacement between consecutive volumes or ≥3 mm total displacement over the course of a 7 min run. After runs that exceeded these thresholds were excluded, two participants had fewer than five remaining runs and so were excluded entirely as there were insufficient data for MVPA. Participants were screened for stereoacuity (<1° arcmin near/far discrimination threshold) and contraindications to MRI prior to the experiment. All experiments were conducted in accordance with the ethical guidelines of the Declaration of Helsinki and were approved by the University of Cambridge STEM and all participants provided informed consent.

### Stimuli: main experiment

To assess whether global context alters local stereoscopic V1 activity, we ran two depth conditions: 1) an ambiguous “wallpaper” condition in which two stereoscopic matches are likely and the perceived depth of the stimulus is disambiguated by the background luminance of the pattern, and 2) a standard disparity condition in which the perceived depth of the stimulus is produced by introducing a position shift between the patterns presented in the left and right eyes. Stimuli were full contrast greyscale stereograms of square-wave gratings, 4.8° wide and 4° high, surrounded by a 0.6° thick light-grey or dark-grey background (**Figure 3**). The spatial period of the square-wave grating was 1.2°, resulting in 8 segments in each pattern.

**Figure 3.**
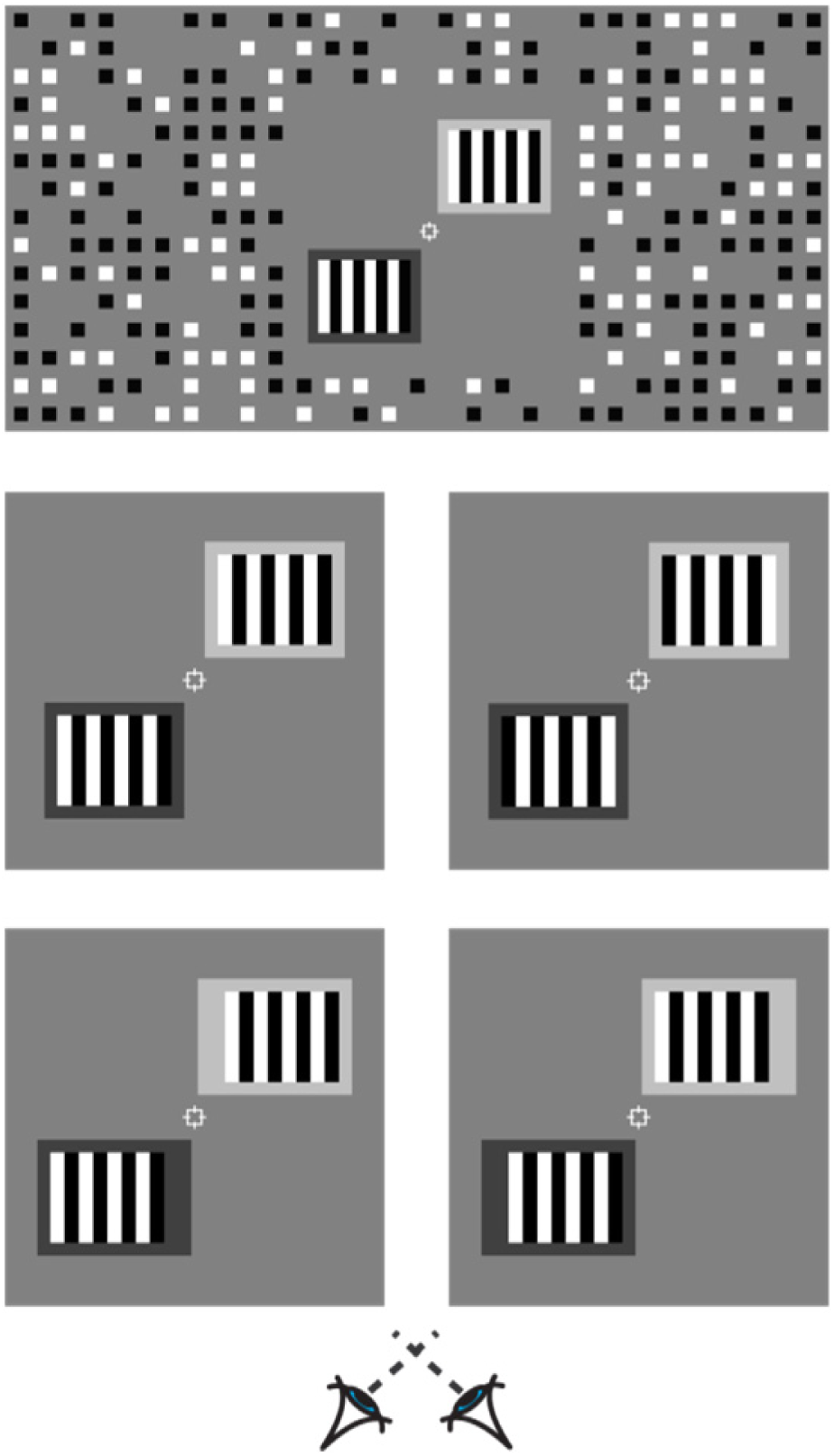
Illustration of the (top) entire stimulus as seen through one eye, (middle) ambiguous depth and (bottom) standard depth stimuli used in the experiment; cross-fuse to observe depth.

In the ambiguous depth condition, a 180° phase offset in the square-wave was introduced between stimuli presented in the left and right eye, producing a ±36 arcmin binocular disparity when fused. The binocular disparity was relatively large, as it is dependent on the spatial period of the grating, and this was required to be sufficiently large to permit effective localization of different regions of the pattern, i.e., the centre and flankers. In the standard depth condition, the phase of the gratings was kept constant between the left and right eyes, while the position of the gratings was shifted 18 arcmin horizontally in each eye, to produce an offset (i.e., either near or far) corresponding to that produced by matching the gratings in the ambiguous condition. (It is worth clarifying that in this standard depth condition, there is an unambiguous overall disparity, but, by design, the central portion of the stimulus contains local information that could be matched in a temporal or a nasal direction for a neuron with a small receptive field. This is common to almost all binocular images containing self-similar elements, and is conceptually similar to the motion aperture problem.) The position and size of the background in the standard depth condition was adjusted so as to appear equivalent to that in the ambiguous condition.

Two stereogram gratings were presented on each trial, in the top-right and bottom-left quadrants of the screen, centred on the oblique axis, at a distance of 4.8° from the centre. In addition to the phase offset between gratings presented to the left and right eye, the top and bottom gratings were offset by 180°. Thus, there were eight conditions: ambiguity (ambiguous/standard) × background in the top/bottom (dark-light/light-dark) × phase in the top/bottom (0°-180°/180°-0°). These conditions were designed such that there were two possible depth configurations (near-far, far-near), equally balanced across conditions (that is, four of each). See **Table 1** for a summary of the experimental manipulations across conditions. Stimuli were presented on a mid-grey background, surrounded by a grid of black and white squares (75% density) designed to provide an unambiguous background reference.

**Table 1.** Summary of experimental manipulations.

| Condition number | Ambiguous | Background luminance in the top/bottom | Pattern phase shift in the top/bottom | Pattern position shift in the top/bottom | Perceived depth in the top/bottom |
| --- | --- | --- | --- | --- | --- |
| 1 | No | Light, dark | 0, 180 | Crossed, uncrossed | Near, far |
| 2 | No | Dark, light | 180, 0 | Crossed, uncrossed | Near, far |
| 3 | No | Light, dark | 180, 0 | Uncrossed, crossed | Far, near |
| 4 | No | Dark, light | 0, 180 | Uncrossed, crossed | Far, near |
| 5 | Yes | Light, dark | 0, 180 | None | Near, far |
| 6 | Yes | Dark, light | 180, 0 | None | Near, far |
| 7 | Yes | Light, dark | 180, 0 | None | Far, near |
| 8 | Yes | Dark, light | 0, 180 | None | Far, near |

### Stimuli: checkerboard localizer

We localized regions of interest (ROIs) corresponding to the spatial location of the experimental stimulus for each participant in two separate runs. Subjects were presented with full contrast flickering checkerboard stimuli (10 Hz, black and white checks, check size = 12 × 12 arcmin; **Figure 4**). The localizer contained four types of trials, corresponding to the locations of ROIs of the experimental stimulus: the central six segments (center) of the (1) left and (2) right patterns, (3) the two outer segments (flankers), and (4) the background. The contrast of the checkerboards decayed according to a cosine function over 12 arcmin from the edges.Stimuli were programmed and presented in MATLAB (The MathWorks, Natick, MA) with Psychophysics Toolbox extensions (Brainard, 1997; Pelli, 1997). Stereoscopic presentation in the scanner was achieved using a “PROPixx” DLP LED projector (VPixx Technologies) with a refresh rate of 120 Hz and resolution of 1920 × 1080, operating in RB3D mode. The left and right images were separated by a fast-switching circular polarization modulator in front of the projector lens (DepthQ; Lightspeed Design). The onset of each orthogonal polarization was synchronized with the video refresh, enabling interleaved rates of 60 Hz for each eye’s image. MR-safe circular polarization filter glasses were worn by subjects in the scanner to dissociate the left and right eye’s view of the image. Stimuli were back-projected onto a polarization-preserving screen (Stewart Filmscreen, model 150) inside the bore of the magnet and viewed via a front-surfaced mirror attached to the head coil and angled at 45° above the observers’ heads. This resulted in a viewing distance of 82 cm, from which both wallpaper stimuli were visible within the binocular field of view.

**Figure 4.**
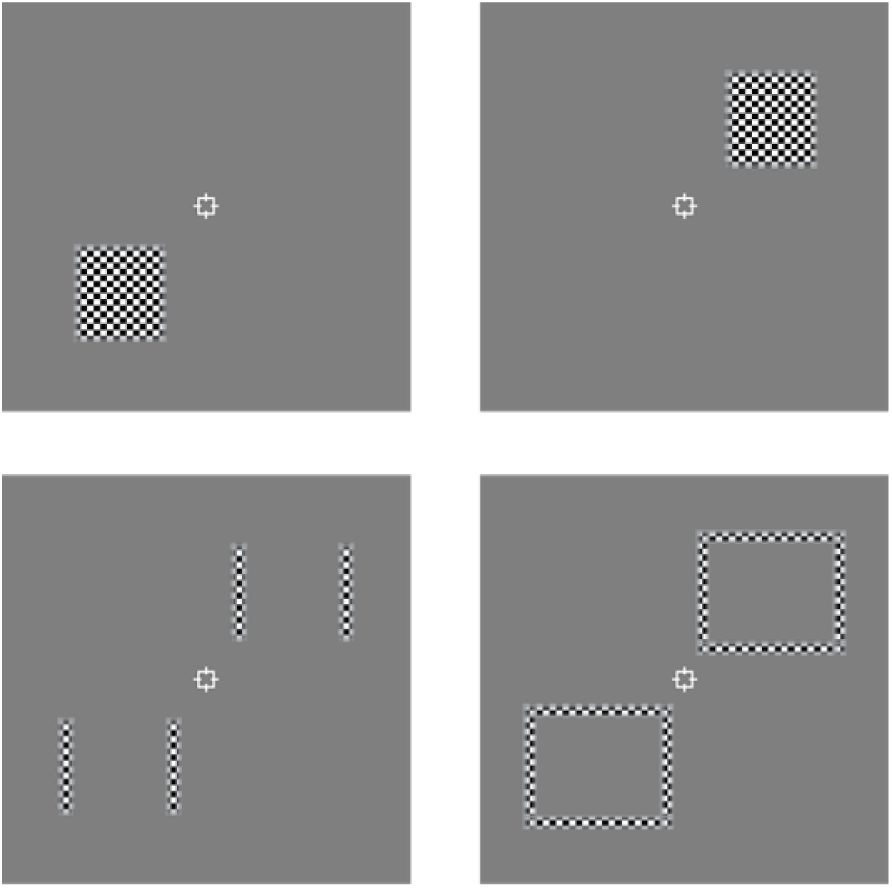
Illustration of the checkerboard stimuli used for functional localization of the (top) left and right centres, (bottom-left) flankers, and (bottom-right) background, of the stimuli.

### Vernier task

During both experimental and checkerboard localizer scans, participants performed an attentionally demanding Vernier task at fixation. Participants were instructed to fixate a central cross hair fixation marker. The fixation marker consisted of a white square outline (side length 30 arcmin) and horizontal and vertical nonius lines (length 22 arcmin). One horizontal and one vertical line were presented to each eye in order to promote stable vergence and to provide a reference for a Vernier task (Popple, Smallman, & Findlay, 1998). The Vernier target line subtended 6.4 arcmin in height by 2.1 arcmin in width and was presented at seven evenly spaced horizontal offsets of between ±6.4 arcmin for 500 ms (with randomized onset relative to stimulus) on 20% of TRs during experimental scans and 50% during checkerboard localizer scans. During the checkerboard localizer scans, participants were instructed to indicate, by button press, which side of the central upper vertical nonius line the target appeared, and the target was presented monocularly to the contralateral eye. During the experimental scans, observers were instructed to indicate, by button press, whether the pattern on the same (left/right) side as the vernier line was near or far, relative to the background. For both scan types the vernier task served to maintain observers’ fixation on the central crosshair, while in the experimental scans it also served to provide a measure of the perceptual strength of depth.

### Psychophysics

In a separate session, prior to being scanned, participants performed a single-interval forced-choice discrimination task in which they were presented one of the four stimuli from the ambiguous depth conditions, drawn pseudo-randomly. Their task was to indicate, via (up/down) button press, the location of the grating that appeared nearer than the background. No duration restriction was enforced, but observers were asked to respond as soon as they made a judgement. Observers completed 50 trials for each condition, for a total of 200 trials. For each condition, responses were assigned either a 1 (up) or −1 (down) and the average was taken to produce a measure of performance.

### Imaging

Data were collected at the Cognition and Brain Sciences Unit with a 3T Siemens Prisma MRI scanner with a 32 channel head coil. Blood oxygen level-dependent (BOLD) fMRI data were acquired with a gradient echo echo-planar imaging (EPI) sequence [echo time (TE) 29 ms; repetition time (TR) 2000 ms; voxel size 1.5 × 1.5 × 2 mm, 30 slices covering occipital cortex] for experimental and localizer scans. Slice orientation was close to coronal section but rotated about the mediolateral axis with the ventral edge of the volume more anterior in order to ensure coverage of lateral occipital cortex (**Figure 5a**). A high-resolution T1-weighted anatomical scan (voxel size 1 mm isotropic) was additionally acquired for each participant.

**Figure 5.**
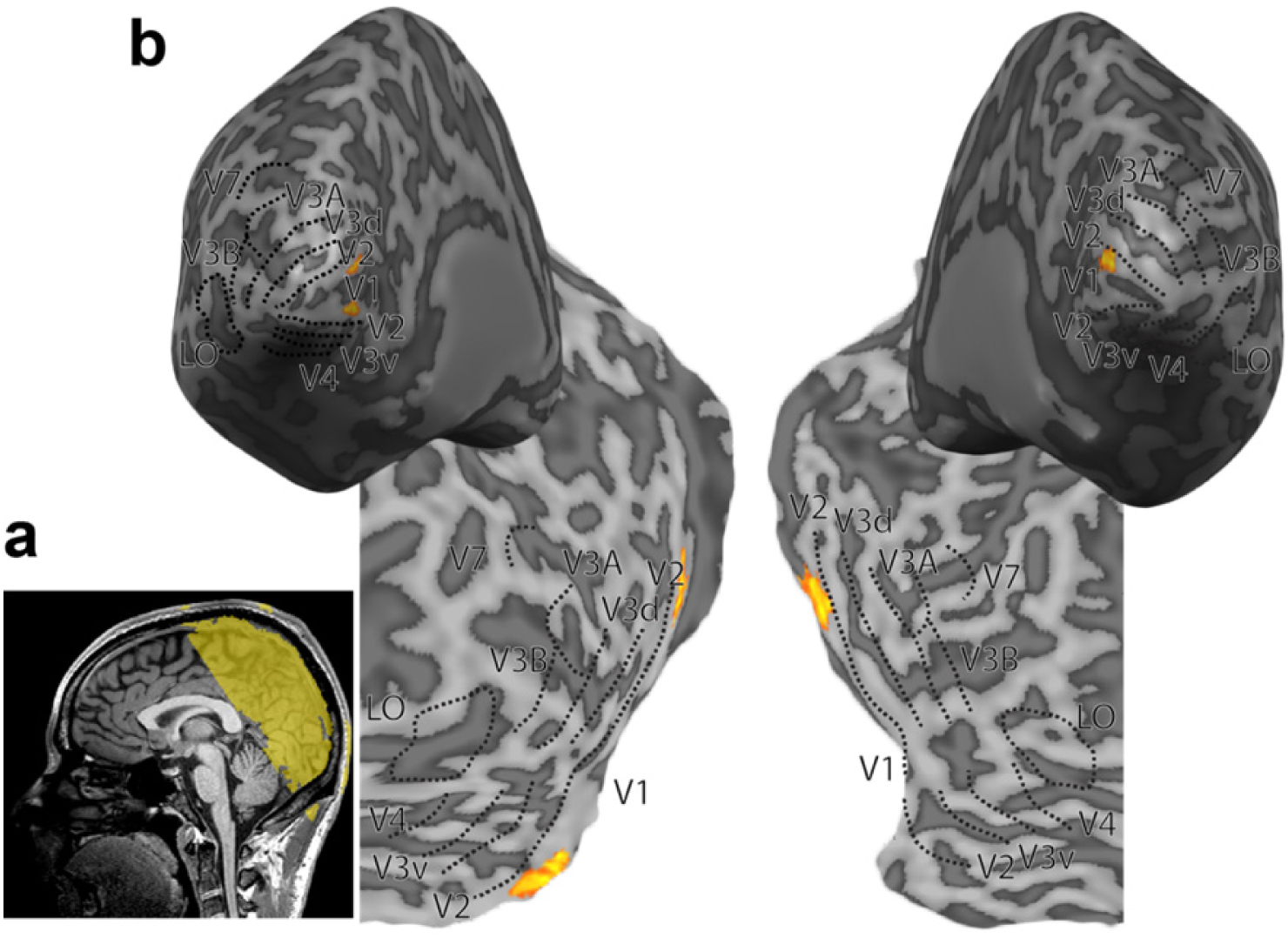
**a)** Representative inflated cortical surface reconstruction and flat maps showing the visual regions of interest from 1 participant. These include retinotopic areas, V3B/KO, and lateral occipital (LO) area. Sulci are rendered a darker grey than gyri. Superimposed on the maps are “centre subROIs”. **b)** Sagittal view of a structural MRI scan of 1 participant, with coloured overlay illustrating slice orientation for functional scans

### Retinotopy

In a separate session, we localized ROIs for each participant using standard retinotopic mapping procedures (**Figure 5b**). Retinotopically organized visual areas V1, V2, V3v, V4, V3d, V3A, V3B/KO, and V7 were defined with polar and eccentricity maps, which were generated by fMRI responses to rotating wedge and expanding concentric ring stimulus presentations, respectively (DeYoe et al., 1996; Sereno et al., 1995). V3 was divided into dorsal and ventral quadrants in each hemisphere (V3d and V3v) in line with previous delineation based on both functional and cytoarchitectonic distinctions (Wilms et al., 2010). Area V4 was defined as the region of ventral visual cortex adjacent to V3v containing a representation of the upper quadrant of the contralateral visual field (Tootell, 2001; Tyler et al., 2005). Area V7 was defined as the region of retinotopic activity anterior and dorsal to V3A (Tootell, 2001; Tsao et al., 2003; Tyler et al., 2005). Area V3B/KO (Tyler, Likova, Kontsevich, & Wade, 2006; Zeki, Perry, & Bartels, 2003) was defined as the area comprising the union of retinotopically mapped V3B and an area “KO” that was functionally localized by its preference for motion-defined boundaries compared with transparent motion of black and white dots (P < 10^4^) (Dupont et al., 1997; Tootell, 2001; Tyler et al., 2006; Zeki et al., 2003). This area contained a full hemifield representation, inferior to V7 and lateral to, and sharing a foveal confluence with, V3A (Tyler et al., 2005). Lateral occipital complex (LOC) was defined as the set of voxels in lateral occipito-temporal cortex that responded significantly more strongly (P < 10^4^) to intact, compared with scrambled, images of objects (Kourtzi, Betts, Sarkheil, & Welchman, 2005). LOC subregion LO, which extended into the posterior inferotemporal sulcus, was defined based on the overlap of functional activations and anatomical structures, consistent with previous studies (Grill-Spector, Kushnir, Hendler, & Malach, 2000).

### fMRI design

In the fMRI experiment, each run consisted of 24 blocks and began and ended with a 16 s fixation period during which only the background and fixation marker were presented. Each block was 16 s in duration. The main experimental blocks consisted of a stimulus from one of the eight conditions being presented for 1 s followed by a 1 s fixation period, thus stimuli were presented for a total of 8 s per 16 s block. The checkerboard blocks were the same (24 blocks), except the stimulus was drawn from one of the four checkerboard conditions. Total block run length was 7 min 4s. Block order was randomized, and each of the eight/four conditions was presented for three blocks per run. Observers completed a total of nine runs: seven of the main experiment and two of the checkerboard localizer (at the beginning and the end).

### fMRI data analysis

Anatomical scans of each participant were transformed into Talairach space (Jean & Tournoux, 1988), and inflated and flattened surfaces were rendered with BrainVoyager QX (BrainInnovation, Maastricht, The Netherlands). Functional data were preprocessed with slice timing correction, head motion correction, and high-pass filtering before being aligned to the participant’s anatomical scan and transformed into Talairach space. Global signal variance across each run was calculated and (2) runs with excessive variance (>.5) were removed.

### ROI analysis

Within each condition, we tested the discriminability of BOLD responses to the two different depth configurations (near-far vs. far-near). For example, for the standard disparity condition we compared the activity of runs from conditions one and two and to three and four (**Table 1**). For each ROI we selected grey matter voxels from both hemispheres and ranked them on the basis of their response (as indicated by *t*-values calculated with the standard general linear model) to a contrast between the “centre” region localizer and fixation. Thus, voxel selection was based on data independent from the classification data and prioritized voxels responding to the region of the stimulus that was perceived as near or far. Assessment of the point at which classification accuracies tended to saturate as a function of pattern size resulted in the selection of 200 voxels per ROI. This voxel selection process has been validated by previous studies (Kamitani & Tong, 2005; Preston, Li, Kourtzi, & Welchman, 2008).

We normalized each voxel time course separately for each experimental run (by calculating z scores) to minimize baseline differences across runs. The multivariate analysis data vectors were produced by applying a 4 s temporal shift to the fMRI time series to compensate for the lag time of the hemodynamic response function and then averaging all time series data points of an experimental condition. We used a support vector machine (SVM; SVM^light^ toolbox, http://svmlight.joachims.org) for classification and performed a sixfold leave-one-out cross-validation, in which data from six runs were used as training patterns (24 patterns, 4 per run for each configuration) and data from the remaining run were used as test patterns (4 per run for each configuration). For each participant, the mean accuracy across cross-validations was taken.

Previous work has shown that depth can be accurately decoded from retinotopic areas in the primary visual cortex (Ban, Preston, Meeson, & Welchman, 2012; Preston et al., 2008). There were three subjects for whom the average decoding accuracy across all retinotopic areas in the standard disparity condition was not significantly above chance. Including data from subjects for whom standard disparity cannot be decoded reduces the interpretability of the results from the ambiguous condition; thus, these subjects were omitted from the analysis. For the remaining ten subjects, the average decoding accuracy was relatively high (63%), indicating that the (3) omitted subjects had poor data quality. We screened observers for discrimination of small disparities (>1 arcmin), whereas the disparity used in the experiment was relatively large (±36 arcmin); thus, one explanation for the inability to decode depth in the standard condition for these three observers is that they suffered from specific stereo deficits for large disparities (Van Ee & Richards, 2002), which we did not detect during screening, and could not perceive the depth of the stimuli. Alternatively, observers may have had specific difficulty in perceiving the depth of the stimuli as they were presented in the fMRI scanner.

### SubROI analysis

Data from the checkerboard localizer scans was averaged over each block to define V1 ‘subROIs’. SubROIs were defined by contrasting each V1 voxel’s response between checkerboard localizer conditions (as indicated by *t*-values calculated with the standard general linear model). For instance, in a contrast between the centre localizer and the flanker and background localizers, voxels with *t*-value larger than 2.0 were considered “centre voxels”. A cluster correction was then used to remove potentially spurious groups of <25 voxels, and the remaining voxels were considered subROIs (**Figure 5b**). This procedure resulted in the selection of 85 voxels per subROI. For some subROIs in some subjects, 85 voxels were not available, in which case we used the maximum number of voxels available.

## RESULTS

### Psychophysics

We measured the perceptual consistency of depth produced by the stimuli in the ambiguous condition for all participants. Specifically, we presented each of the four (background × phase) stimuli and assessed the frequency with which participants reported the same depth configuration (near-far or far-near). A score of one indicates the participant perceived a near-far configuration on all trials, a score of minus one indicates they perceived a far-near configuration on all trials, and a score of zero indicates they perceived it equally often in each configuration. The ideal observer would always perceive the stimuli in a manner where binocular matches preserve contrast polarity (**Figure 6a**). All participants showed the same pattern of “polarity preserving” results, with varying degrees of consistency. For each condition, group average performance was significantly different than zero (**Figure 6b**).

**Figure 6.**
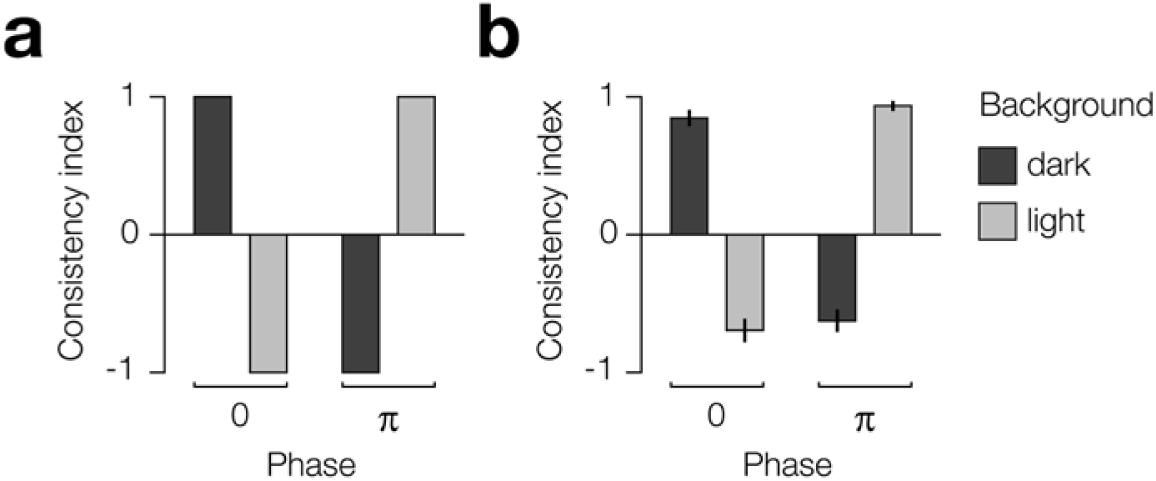
Predicted ideal observer and actual results of psychophysical test of ambiguous wallpaper task. **a)** Predicted depth judgements in each of the four (background × phase) conditions based on binocular matching that preserves contrast polarity. **b)** Average group performance across conditions. The consistency index indicates the proportion of depth configuration judgements that were near-far: 1 = 100%, 0 = 50%, and −1 = 0%.

### fMRI

To assess information about the displayed stimulus contained in the fMRI response, we used a machine learning classifier to decode the stimulus from the brain activity. We did this across the multiple ROIs we had sampled, for both the ambiguous and standard stimuli (**Figure 7a**). A repeated measures analysis of variance (ANOVA) revealed a main effect of condition (Ambigous vs. standard depth; *F*_1,9_=26.1, *P*=6.4e^-4^) but not ROI (*F*_8,72_=1.2, *P*=.28). Consistent with previous work (Ban et al., 2012; Preston et al., 2008), we were able to decode depth from (standard) binocular disparity across all of the retinotopic areas we sampled (*F*_1,9_=47.9, *P*=6.9e^-5^), with highest accuracies found in visual area V3A (**Figure 7a**). By contrast, when we trained the classifier on fMRI activity evoked by ambiguous stimuli, we were unable to decode depth reliably in any of the ROIs (*F*_1,9_=0.001, *P*=0.97). A possible explanation for this is that the percept of depth was less robust for ambiguous stimuli, evoking more variable fMRI responses that provided a poor basis on which to train the machine learning classifier. We therefore tested whether depth could be decoded for ambiguous stimuli when the classifier had been trained on fMRI activity evoked by the standard stimuli and vice versa (**Figure 7b**). As we expected, we found that we were unable to reliably decode standard depth using a classifier trained on ambiguous depth (*F*_1,9_=3.5, *P*=0.20), supporting the interpretation that the ambiguous stimuli provided a poorer training set than the standard stimuli. Similarly, were we unable to decode ambiguous depth using a classifier trained on activity evoked by the standard the depth stimuli (*F*_1,9_=3.5, *P*=0.09).

**Figure 7.**
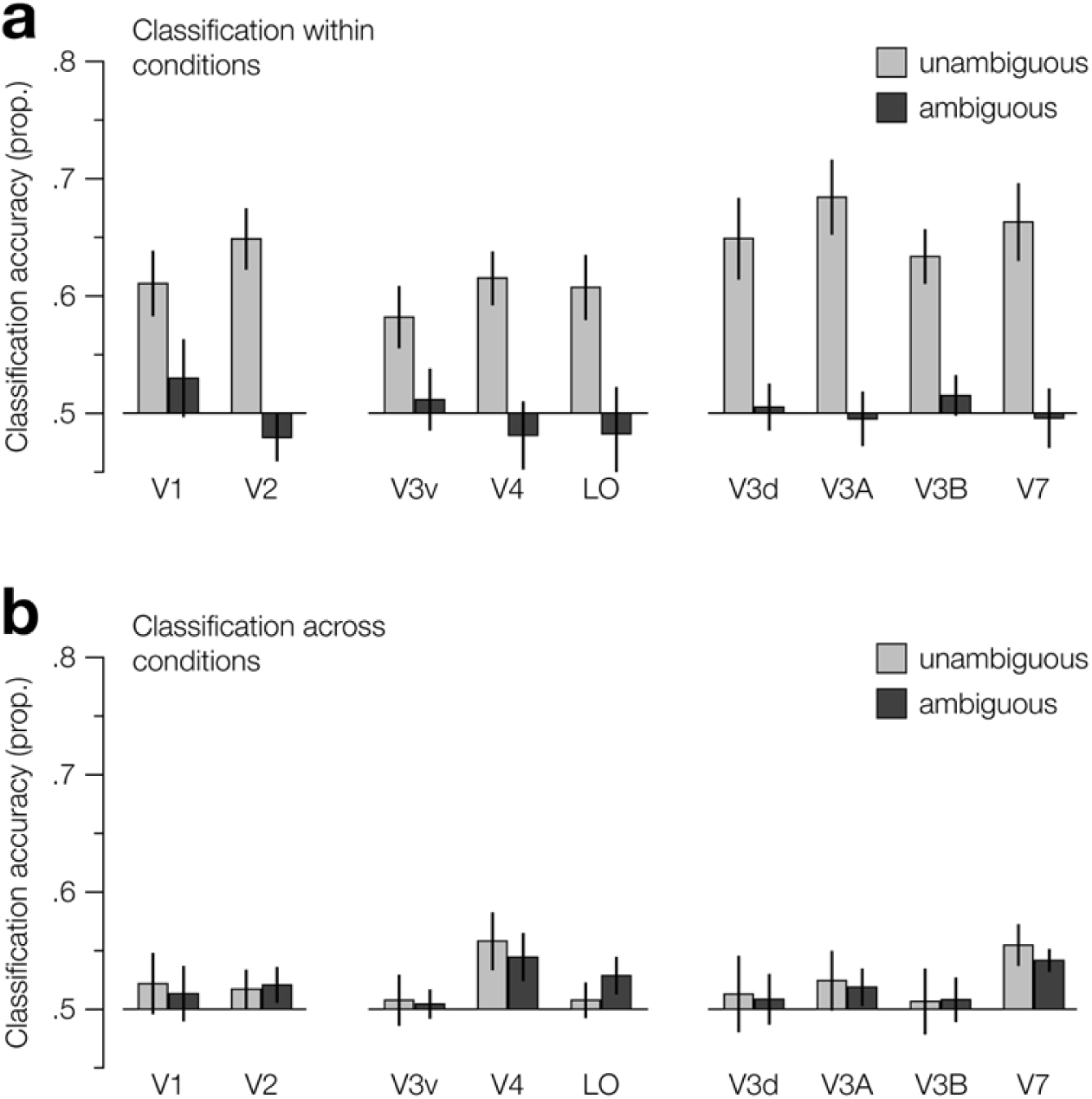
Classifier decoding accuracy for discriminating far-near vs. near-far depth stimulus configurations, for all retinotopic regions of interest. Prediction performance is shown for (a) classifiers trained and tested on activity within ambiguity conditions and (b) classifiers trained on activity from one ambiguity condition and tested on the other (transfer test). For (b), the legend labels refer to the condition that the classifiers were tested on. For all regions, we used performance based on activity of 200 voxels. Error bars indicate s.e.m.

When we considered the aggregate activity of voxels within area V1, we found that we were unable to reliably decode depth from fMRI activity evoked by ambiguous stimuli (**Figure 7**). However, our particular interest was to examine responses within subregions of the viewed stimuli. Thus, to test whether global context influences local stereoscopic-related activity in V1 we used MVPA to decode fMRI activity from each of the subROIs corresponding to the centre, flankers, and background of the stimulus. If V1 activity is influenced by global context, we would expect to be able to decode depth from activity in the flankers and centre subROIs, but not the background. That is, the background should influence the perceived depth of the pattern, but not the other way around.

We first tested whether we could reliably decode the depth of standard stimuli from the fMRI activity within the V1 subROIs. While the central region of the standard stimulus could also provide ambiguous information to neurons with spatially-limited receptive fields, binocular matches are possible for all elements of the stereogram, i.e., there is a single probable matching solution that is unaffected by the luminance of the background. Thus, while decoding activity evoked by standard stimuli is not diagnostic of contextual influence, it provides a sanity check that perceived depth can be decoded from the activity in these subregions. We found that we could reliably decode depth from all subROIs (centre, *t*_9_=5.18, *P*=5.8e^-4^; flankers, *t*_9_=4.21, *P*=.002; background, *t*_9_=4.41, *P*=.002; **Figure 8a**). This would appear to be contradictory of the previous prediction, i.e., activity from the background should be undiagnostic of depth; however, the subROI localizers were designed for the ambiguous stimuli, where the position of the pattern is held constant. In contrast, we shifted the position of the standard stimuli nasally and temporally to produce a binocular disparity, meaning that the areas contained by the background subROI also contained partial flanker sections of these stimuli. These monocular cues, i.e., whether the pattern was shifted nasally or temporally, explain why we were still able to reliably decode the depth configuration from activity within the background subROI.

**Figure 8.**
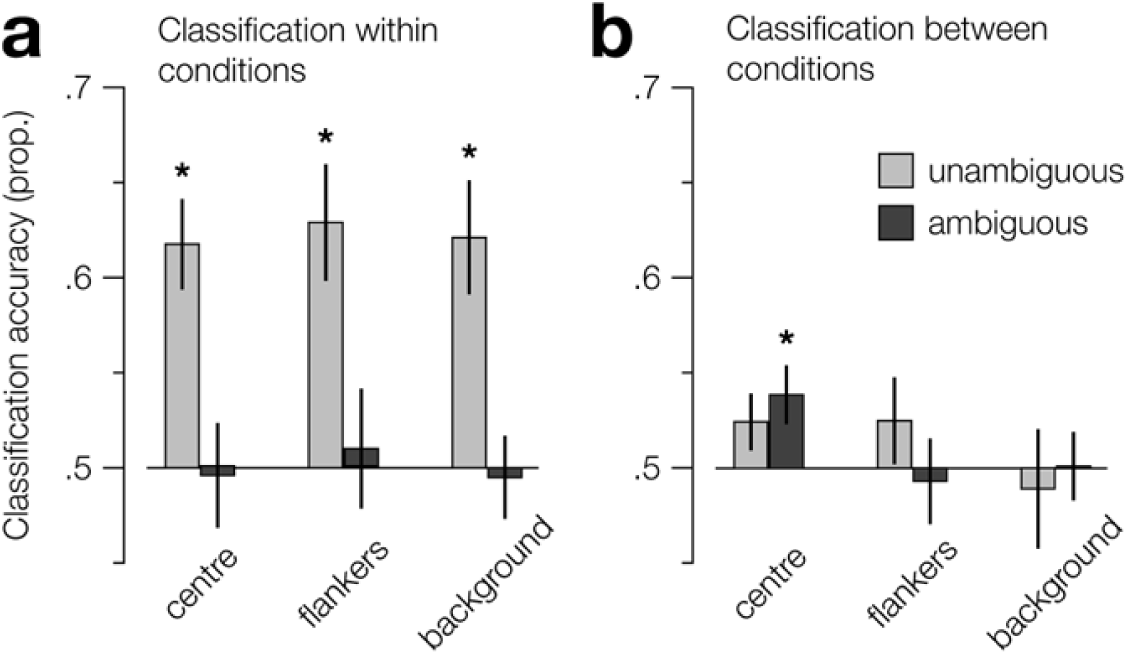
Classifier decoding accuracy for discriminating far-near vs. near-far depth stimulus configurations, for stimulus-related V1 subregions of interest. Prediction performance is shown for (a) classifiers trained and tested on activity within ambiguity conditions and (b) classifiers trained on activity from one ambiguity condition and tested on the other (transfer test). For (b), the legend labels refer to the condition that the classifiers were trained on. For all regions, we used performance based on activity of 85 voxels. Error bars indicate s.e.m. Asterisks indicate areas where performance is significantly above chance.

We next tested whether we could decode the depth of the ambiguous stimuli from the activity within V1 subROIs. Consistent with the retinotopic ROI analysis, we were unable to reliably decode depth of ambiguous stimuli from any of the subROIs using a classifier trained on the ambiguous depth stimuli (**Figure 8a**). We then tested for transfer between the conditions by training the classifier on fMRI activity evoked by stimuli in one condition and testing on activity evoked by stimuli in the other. Using a classifier trained on standard depth stimuli evoked activity, we were able to reliably decode the depth of the ambiguous stimuli from activity in the centre subROI (*t*_9_=2.58, *P*=.029; **Figure 8b**). As the input to this subregion was ambiguous, i.e., information from the background/edges is needed to disambiguate the signal, the fact that depth can be reliably decoded from its activity shows that it is influenced by the global context of the stimulus, i.e., the edges/background. Consistent with the results from the ROI analysis and the interpretation that the ambiguous dataset could not be used to train an effective classifier, using a classifier trained on activity evoked by ambiguous stimuli, we were unable to reliably decode the depth of the standard stimuli using from activity from in the same (centre) subregion (*t*_9_=1.63, *P*=.138).

We were unable to decode depth form the flankers and background (**Figure 8b**). As noted previously, to produce the percept of depth for the standard stimuli the position of the pattern was shifted horizontally between left and right eyes, resulting in misalignment of the flankers and background relative to the subregion localizers. To decode the “depth” in the flanker and background subregions, the classifier could rely on monocular cues, i.e., the relative position of the flankers to the background. By contrast, these monocular cues were not present in the ambiguous stimuli, as the position of the pattern was held constant. Thus, it is likely that training did not transfer from standard to ambiguous stimuli in these subregions as the classifier could not use the same monocular cues to decode fMRI activity from the ambiguous test set.

We designed the experiment such that the stimulus manipulations were balanced across conditions. This allowed us to test the stimulus information that the classifier was capable of decoding from the fMRI activity within each subregion. We carefully designed the localizers to only include voxels whose activity was evoked by specific subregions of the ambiguous stimulus and contrasted between them to exclude shared voxels. However, a possible concern is that some of the voxels may have sampled information from the other subregions as a result of eye movements. Thus, we tested from which subregions the (light/dark) luminance of the background could be reliably decoded. As expected, we found that background luminance could be reliably decoded from the background subROI (*t*_9_=3.04, *P*=.014). Critically, we were not able to reliably decode the background luminance of the ambiguous stimuli from the centre subROI using a classifier trained on either activity evoked from ambiguous (*t*_9_=1.53, *P*=.16) or standard stimuli (*t*_9_=1.03, *P*=.33). This shows that the capacity to decode depth from (ambiguous stimuli evoked) activity in the centre subregion was not due to unintentional sampling from the background, but evidence that stereoscopic-related activity in V1 is influenced by global context.

Another possible concern is that decoding accuracy may reflect eye movements made in response to stimulus presentation. However, this seems unlikely for two reasons. First, observers were required to perform an attentionally demanding Vernier task, which required constant monitoring of the fixation. Second, the depth of stimuli was balanced within presentations, i.e., each presentation consisted of a near and a far stimulus, thus there was no global difference in depth between conditions to promote one pattern of vergence eye movements over another.

## DISCUSSION

The offset between an image projected onto the left and right retinae (binocular disparity) is sufficient to produce an estimate of depth. To compute this offset, it has long been understood that the visual system must first match the features of the image between the retinae (“solve the correspondence problem”). This problem is nontrivial, as the number possible solutions rapidly increases with the number of to-be-matched elements. Classically, investigations of the correspondence problem in stereopsis have focused on the how local features facilitate matching. In contrast, with the exception of a few notable studies (Anderson & Nakayama, 1994; McKee, Verghese, Ma-Wyatt, & Petrov, 2007), the role of global context has not been tested extensively.

To test whether global context influences the activity of V1 during binocular matching, here we used fMRI to measure observers’ brain activity while viewing ambiguous stereograms, whose binocular matching (producing the perception of either near or far depths) was determined by the luminance of the background. First, we used multi-voxel pattern analysis (MVPA) of the fMRI activity to decode depth from retinotopically defined ROIs (V1, V2, V3d, V3A, V3B/KO V4, V7, & LO). Consistent with previous work (Ban et al., 2012; Preston et al., 2008), for standard disparity-defined stimuli we find that depth can be reliably decoded from all ROIs, with maximum decoding accuracy in V3A. By contrast, for ambiguous stimuli, we find that depth cannot be reliably decoded from any of the ROIs we tested. Next, we tested whether depth could be decoded from the activity of V1 subROIs receiving ambiguous binocular input while perception is influenced by the surrounding background. We found that near vs. far depth could be reliably decoded from regions of V1 that only receive ambiguous information, indicating that the local activity of V1 is influenced by global context.

While we were able to reliably decode depth from activity evoked by the centre subregion of the ambiguous stimulus using a classifier trained on the standard depth stimuli, we were unable to decode depth using a classifier trained on ambiguous stimuli. This is most probably explained by the ambiguous stimulus providing a less stable interpretation of depth: while manipulating the background luminance biases perceived depth, there remains some bistability in the percept. Thus, for some presentations of the ambiguous stimulus, observers may have perceived the opposite depth to that which the activity was labelled, reducing the capacity of the SVM to learn a pattern of activity associated with the label. Despite this, it is reassuring to find evidence of transfer of decoding accuracy for a classifier trained on standard depth signals and tested on ambiguous depth signals, as it indicates that the SVM is using (at least partially) shared stereoscopic-related brain activity to classify the stimuli.

The finding that V1 stereoscopic-related BOLD activity is influenced by context seems at odds with previous neurophysiological work indicating the activity of disparity-selective V1 neurons does not reflect depth perception (Cumming & Parker, 2000, 1997). However, for many neurons the activity produced by anti-correlated matches is attenuated compared to that produced by correlated matches (Cumming & Parker, 1997). Thus, a possible mechanism of the global influence on V1 activity observed here may be through inhibitory feedback or lateral connections, perhaps originating from a higher area involved in resolving perceptual ambiguity.

A limitation of the current study is that it cannot distinguish the two possible mechanisms through which contextual influence in V1 is realized, that is, lateral and feedback connections. Previous theoretical modelling (Friston, 2005; Mumford, 1992; Phillips, Clark, & Silverstein, 2015) and fMRI work investigating other effects of spatial context in vision (Kok & De Lange, 2014; Muckli et al., 2015) suggest that contextual effects are facilitated by feedback, rather than lateral, connections within the primary visual cortex; however, to resolve this ambiguity for the effect of global context on stereoscopic activity, future research could employ fMRI with higher spatial resolution to examine activity at different laminar layers (Muckli et al., 2015).

In sum, here we provide evidence that the global context in stereoscopic matching can modulate the activity of binocular stimuli in the primary visual cortex. These findings have implications for models of stereopsis, which focus on local binocular matching, underscoring the importance of considering both local and global stereoscopic activity, as early as V1, in the computation of depth from binocular disparity.

## ACKNOWLEDGEMENTS

This work was supported by the Leverhulme Trust (ECF-2017-573 to RR) and the Wellcome Trust (095183/Z/10/Z to AEW).

## REFERENCES

Anderson, B. L., & Nakayama, K. (1994). Toward a general theory of stereopsis: Binocular matching, occluding contours, and fusion. Psychological Review, 101(3), 414–445. http://doi.org/10.1037/0033-295X.101.3.414

Ban, H., Preston, T. J., Meeson, A., & Welchman, A. E. (2012). The integration of motion and disparity cues to depth in dorsal visual cortex. Nature Neuroscience, 15(4), 636–643. http://doi.org/10.1038/nn.3046

Barlow, H. B., Blakemore, C., & Pettigrew, J. D. (1967). The neural mechanism of binocular depth discrimination. The Journal of Physiology, 193(2), 327–342. http://doi.org/10.1113/jphysiol.1967.sp008360

Brainard, D. H. (1997). The Psychophysics Toolbox. Spatial Vision, 10(4), 433–436. http://doi.org/10.1163/156856897X00357

Brewster, D. (1844). XLIII.— On the Knowledge of Distance given by Binocular Vision. Transactions of the Royal Society of Edinburgh, 15(4), 663–675. http://doi.org/10.1017/S0080456800030246

Cumming, B. G., & Parker, a J. (2000). Local disparity not perceived depth is signaled by binocular neurons in cortical area V1 of the Macaque. The Journal of Neuroscience?: The Official Journal of the Society for Neuroscience, 20(12), 4758–4767. http://doi.org/20/12/4758[pii]

Cumming, B. G., & Parker, A. J. (1997). Responses of primary visual cortical neurons to binocular disparity without depth perception. Nature, 389(6648), 280–283. http://doi.org/10.1038/38487

DeYoe, E. A., Carman, G. J., Bandettini, P., Glickman, S., Wieser, J., Cox, R., … Neitz, J. (1996). Mapping striate and extrastriate visual areas in human cerebral cortex. Proceedings of the National Academy of Sciences of the United States of America, 93(6), 2382–6. http://doi.org/10.1073/pnas.93.6.2382

Dupont, P., De Bruyn, B., Vandenberghe, R., Rosier, A.-M., Michiels, J., Marchal, G., … Orban, G. A. (1997). The kinetic occipital region in human visual cortex. Cerebral Cortex, 7(3), 283–292. http://doi.org/10.1093/cercor/7.3.283

Friston, K. (2005). A theory of cortical responses. Philosophical Transactions of the Royal Society B: Biological Sciences, 360(1456), 815–836. http://doi.org/10.1098/rstb.2005.1622

Grill-Spector, K., Kushnir, T., Hendler, T., & Malach, R. (2000). The dynamics of object-selective activation correlate with recognition performance in humans. Nature Neuroscience, 3(8), 837–893. http://doi.org/10.1038/77754

Hubel, D. H., & Wiesel, T. N. (1970). Stereoscopic vision in macaque monkey: Cells sensitive to binocular depth in area 18 of the macaque monkey cortex. Nature, 225(5227), 41–42. http://doi.org/10.1038/225041a0

Jean, T., & Tournoux, P. (1988). Co-Planar Stereotaxic Atlas of the Human Brain: 3-D Proportional System: An Approach to Cerebral Imaging (Thieme Classics): J. Talairach: 9780865772939: Amazon.com: Books. Thieme, 192. Retrieved from http://www.amazon.com/Co-Planar-Stereotaxic-Atlas-Human-Brain/dp/0865772932

Kamitani, Y., & Tong, F. (2005). Decoding the visual and subjective contents of the human brain. Nature Neuroscience, 8(5), 679–685. http://doi.org/10.1038/nn1444

Kok, P., & De Lange, F. P. (2014). Shape perception simultaneously up- and downregulates neural activity in the primary visual cortex. Current Biology, 24(13), 1531–1535. http://doi.org/10.1016/j.cub.2014.05.042

Kourtzi, Z., Betts, L. R., Sarkheil, P., & Welchman, A. E. (2005). Distributed neural plasticity for shape learning in the human visual cortex. PLoS Biology, 3(7), 1317–1327. http://doi.org/10.1371/journal.pbio.0030204

McKee, S. P., Verghese, P., Ma-Wyatt, A., & Petrov, Y. (2007). The wallpaper illusion explained. Journal of Vision, 7(14), 10.1–11. http://doi.org/10.1167/7.14.10

Muckli, L., De Martino, F., Vizioli, L., Petro, L. S., Smith, F. W., Ugurbil, K., … Yacoub, E. (2015). Contextual Feedback to Superficial Layers of V1. Current Biology, 25(20), 2690–2695. http://doi.org/10.1016/j.cub.2015.08.057

Mumford, D. (1992). On the computational architecture of the neocortex - II The role of cortico-cortical loops. Biological Cybernetics, 66(3), 241–251. http://doi.org/10.1007/BF00198477

Pelli, D. G. (1997). The VideoToolbox software for visual psychophysics: Transforming numbers into movies. Spatial Vision. http://doi.org/10.1163/156856897X00366

Phillips, W. A., Clark, A., & Silverstein, S. M. (2015). On the functions, mechanisms, and malfunctions of intracortical contextual modulation. Neuroscience and Biobehavioral Reviews. http://doi.org/10.1016/j.neubiorev.2015.02.010

Popple, A. V., Smallman, H. S., & Findlay, J. M. (1998). The area of spatial integration for initial horizontal disparity vergence. Vision Research, 38(2), 319–326. http://doi.org/10.1016/S0042-6989(97)00166-1

Preston, T. J., Li, S., Kourtzi, Z., & Welchman, A. E. (2008). Multivoxel Pattern Selectivity for Perceptually Relevant Binocular Disparities in the Human Brain. Journal of Neuroscience, 28(44), 11315–11327. http://doi.org/10.1523/JNEUROSCI.2728-08.2008

Sereno, M. I., Dale, A. M., Reppas, J. B., Kwong, K. K., Belliveau, J. W., Brady, T. J., … Tootell, R. B. H. (1995). Borders of multiple visual areas in humans revealed by functional magnetic resonance imaging. Science, 268(5212), 889–893. http://doi.org/10.1126/science.7754376

Tootell, R. B. H. (2001). Where is “Dorsal V4” in Human Visual Cortex? Retinotopic, Topographic and Functional Evidence. Cerebral Cortex, 11(4), 298–311. http://doi.org/10.1093/cercor/11.4.298

Tsao, D. Y., Vanduffel, W., Sasaki, Y., Fize, D., Knutsen, T. A., Mandeville, J. B., … Tootell, R. B. H. (2003). Stereopsis activates V3A and caudal intraparietal areas in macaques and humans. Neuron, 39(3), 555–568. http://doi.org/10.1016/S0896-6273(03)00459-8

Tyler, C. W., Likova, L. T., Chen, C.-C., Kontsevich, L. L., Schira, M. M., & Wade, A. R. (2005). Extended Concepts of Occipital Retinotopy. Current Medical Imaging Reviews, 1, 319–329. http://doi.org/10.2174/157340505774574772

Tyler, C. W., Likova, L. T., Kontsevich, L. L., & Wade, A. R. (2006). The specificity of cortical region KO to depth structure. NeuroImage, 30(1), 228–238. http://doi.org/10.1016/j.neuroimage.2005.09.067

Van Ee, R., & Richards, W. (2002). A planar and a volumetric test for stereoanomaly. Perception, 31(1), 51–64. http://doi.org/10.1068/p3303

Wilms, M., Eickhoff, S. B., Hömke, L., Rottschy, C., Kujovic, M., Amunts, K., & Fink, G. R. (2010). Comparison of functional and cytoarchitectonic maps of human visual areas V1, V2, V3d, V3v, and V4(v). NeuroImage, 49(2), 1171–1179. http://doi.org/10.1016/j.neuroimage.2009.09.063

Zeki, S., Perry, R. J., & Bartels, A. (2003). The processing of kinetic contours in the brain. Cerebral Cortex, 13(2), 189–202. http://doi.org/10.1093/cercor/13.2.189

